# Identification of novel protective loci for executive function using the trail making test part B in the Long Life Family Study

**DOI:** 10.1101/2025.07.08.663708

**Authors:** Lihua Wang, Katherine Tanner, Stacy L. Andersen, Stephanie Cosentino, Vaha Akbary Moghaddam, E. Warwick Daw, Jason A. Anema, Shiow Jiuan Lin, Acharya Sandeep, Michael Province, Mary K. Wojczynski

## Abstract

The Trail Making Test (TMT) Part B (TMT-B), a well-established assessment of cognitive function, is a frequent component of diagnostic assessments for Mild Cognitive Impairment and dementia in older adults. Identifying the genetic variants associated with the TMT-B will not only gain insights of genetic determinants of cognitive function, but also the molecular mechanisms for dementia. Published GWAS to date for TMT-B suffer from relatively low power due to the use of population level data and imputation methods. To address these deficits, we used a family-based study design to identify the genetic variants associated with the TMT-B incorporating both genome-wide linkage analysis (GWLS) and whole genome sequencing (WGS). As such, we examined the sequenced genetic determinants of TMT-B using GWLS in over 2000 participants from Long Life Family Study (LLFS). In GWLS, the estimated heritability of TMT-B was 0.29. We detected one significant linkage peak at 15q25 (LOD>3.0). Statistical fine-mapping nominated five variants including three SNPs (*NTRK3*-rs74031103, protective *CEMIP*-rs2271159, and protective *AGBL1*-rs4134376) and two INDELs (protective *KLHL25*-15:85882445:IND, and protective *CEMIP*-15:80893381:IND) contributing to the linkage peak. Four out of these five variants are protective for TMT-B. The rs2271159 SNP influences *CEMIP* expression in cerebellum and hippocampus, while the 15:80893381:IND modulates *CEMIP* expression in blood. Additionally, the variant rs4134376 is a basal ganglia-specific eQTL for *AGBL1*. In conclusion, we utilized GWLS, leveraged multi-omics data (whole genome sequence genomic data, transcriptomic data, and lipidomic data), and identified novel protective variants and genes for TMT-B performance.

## Introduction

The Trail Making Test (TMT), one component of the Halstead-Reitan Neuropsychological Test Battery (HRNB), is commonly used to assess attention, visual scanning, processing speed, and elements of executive function including working memory and complex set maintenance in individuals 15 to 93 years old (1–4). This test is widely use in diagnostic assessments for mild cognitive impairment (MCI) and dementia in older adults, including the National Alzheimer’s Coordinating Center (NACC) Uniform Data Set administered at Alzheimer’s Disease Research Centers throughout the United States.

TMT is divided into two parts, including Part A (TMT-A), a measure of attention, visual scanning and processing speed, involving sequentially connecting 25 circles that are randomly arranged and labeled with numbers 1 to 25. The score is the number of seconds it takes to finish this task. For TMT Part B (TMT-B), 25 circles are randomly arranged and labelled using numbers 1 to 13 and letters A through L. The score is the number of seconds used to sequentially connect the circles, in order, alternating between numbers and letters. Given normal performance on TMT-A, deficits on TMT-B can be attributed to cognitive rather than motor or visual scanning deficits. TMT-B is sensitive to a range of neurologic conditions including MCI and Alzheimer’s disease (AD)(5). In fact, a recent study found that TMT-B may be useful as a screening instrument for preclinical AD(6).

TMT-B may be sensitive to early AD due to its multi-dimensional nature. Not only does it have a strong executive component, but it also places demands on spatial navigation, one of the earliest features of AD (7, 8). Indeed, TMT-B has been associated with various aspects of spatial navigation in everyday life including walking and driving(9, 10), as well as at-risk driving behaviors, and increased collisions(1, 11). This study investigates the genetic contributions to TMT-B performance given its sensitivity to various types of cognitive impairment(12) and its association with impairments in everyday activities. These multiple lines of evidence indicate there may be shared brain pathological alterations and genetic architectures underlying TMT-B score and AD. TMT-B has a relative high estimated heritability (between 0.22 and 0.65)(13–16). The investigation of genetic contribution to TMT-B score, may shed light not only on cognitive impairment, but also on the molecular mechanisms of AD. Yet, current understanding of genetic regulation of TMT-B score is limited. To date, the largest GWAS study of TMT for British ancestry (23,822 UK Biobank participants for TMT-A; 23,812 UK Biobank participants for TMT-B) was conducted by Hagenaars SP et al in 2018(16). In this study, genotypes were obtained by imputing variants assayed on the UK BiLEVE array and the UK Biobank axiom array to a UK Biobank-1000 Genomes Phase 3 merged reference panel. Gene-based analyses identified three genes including *CRNKL1*, *CASP5*, and *NAA20* for TMT-A. For TMT-B, one SNP on chromosome 1 (rs34804445), and one gene on chromosome 1 (*PUM1*) reached genome-wide significance (P<5×10^-8^). However, first, the large scale UK Biobank data used by this study is not representative of the general population in US(17). Therefore, variants influencing TMT-B in US population might not be captured by this study. Second, imputation based genetic variants had limited detecting power for rare variants. Third, this study didn’t utilize multi-omics data to map identified variants to functional genes.

To address these limitations, we performed genetic study of TMT-B using the NIA Long Life Family Study (LLFS). LLFS recruited exceptionally long-lived families from US and Denmark. LLFS cohort exhibits a notably healthier aging phenotype compared to participants in the Framingham Heart Study, as evidenced by superior performance across multiple health domains including cognitive function, metabolic health, and physical resilience(18). The availability of well-characterized cognitive traits, including TMT-B, along with various omic data including whole genome sequence data in the LLFS, enables us to investigate the genetic regulatory mechanisms underlying TMT-B performance in long-lived families. Our goal in this study is to assess the genetic determinants of TMT-B performance in exceptionally healthy participants of LLFS using whole genome sequence data, and to map the identified variants to functional molecules by integrating transcriptomic data, lipidomic data, and various QTL resources.

## Methods and Materials

### Samples characteristics

The LLFS is a unique longitudinal, multicenter, and multigenerational family study, aiming to investigate the genetic and environmental determinants of Health Aging Phenotypes (HAPs) and Exceptional Longevity (EL)(18). The LLFS recruited long-lived probands and their family members from four field centers (Boston University School of Medicine in Boston (MA), Columbia College of Physicians and Surgeons in New York (NY), the University of Pittsburgh in Pittsburgh (PA), and the University of Southern Denmark). In LLFS, Baseline (i.e., Visit 1) in-person evaluations (2006–2009) collected various demographic, anthropometric, ankle-brachial index, blood pressure, physical performance, and cognitive function. Visit 2 (2014–2017) gathered the same measures as Visit 1, additional cognitive testing (TMT score), and carotid ultrasonography.. In LLFS, TMT-B was assessed using an Anoto Live Ballpoint Pen (Model DP-20) during the second in-person visit(19). Our analyses utilized 2115 Visit 2 participants who had valid TMT-B score, education (Yes or No for college education), and whole genome sequencing (WGS) data (Table 1).

**Table 1.** Characteristics of samples included in the study.

|  |  |  |  |  |  | Table 1. Characteristics of samples included in the study |  |  |  |  |  |  |  |  |  |
| --- | --- | --- | --- | --- | --- | --- | --- | --- | --- | --- | --- | --- | --- | --- | --- |
| All Families |  |  |  |  |  | Linked Families |  |  |  |  | Top Linked Families |  |  |  |  |
|  | All | Boston | Columbia | Denmark | Pittsburgh | All | Boston | Columbia | Denmark | Pittsburgh | All | Boston | Columbia | Denmark | Pittsburgh |
| N | 2115 | 565 | 365 | 678 | 507 | 397 | 124 | 57 | 112 | 104 | 118 | 51 | 9 | 37 | 21 |
| Female (%) | 1139 (53.85) | 295 (52.21) | 204 (55.89) | 350 (51.62) | 290 (57.20) | 209 (52.64) | 65 (52.42) | 30 (52.63) | 60 (53.57) | 54 (51.92) | 70 (59.32) | 33 (64.71) | 5 (55.56) | 21 (56.76) | 11 (52.38) |
| colledge (%) | 1830 (86.52) | 522 (92.39) | 329 (90.14) | 554 (81.71) | 425 (83.83) | 337 (84.89) | 111 (89.52) | 52 (91.23) | 87 (77.68) | 87 (83.65) | 99 (83.90) | 43 (84.31) | 7 (77.78) | 30 (81.08) | 19 (90.48) |
| Age (avg) | 69.97 (9.97) | 68.46 (8.88) | 70.52 (11.17) | 69.90 (8.60) | 71.37 (11.58) | 69.64 (10.00) | 67.75 (9.29) | 70.77 (10.67) | 71.10 (8.99) | 69.71 (11.20) | 68.64 (10.51) | 68.10 (9.5) | 68.56 (12.33) | 69.86 (10.98) | 67.86 (11.77) |
| AD (%) | 51 (3.55) | 12 (2.12) | 13 (3.56) | NA | 26 (5.13) | 9 (3.16) | 1 (0.81) | 1 (1.75) | NA | 7 (6.73) | 1 (1.23) | 1 (1.96) | 0 (0.00) | NA | 0 (0.00) |
| MCI (%) | 114 (7.93) | 30 (5.31) | 42 (11.51) | NA | 42 (8.28) | 23 (8.07) | 11 (8.87) | 5 (8.77) | NA | 7 (6.73) | 5 (6.17) | 4 (7.84) | 0 (0.00) | NA | 1 (4.76) |
| APOE2 (%) | 372 (17.59) | 98 (17.35) | 71 (19.45) | 122 (17.99) | 81 (15.98) | 77 (19.40) | 26 (20.97) | 12 (21.05) | 25 (22.32) | 14 (13.46) | 26 (22.03) | 14 (27.45) | 3 (33.33) | 7 (18.92) | 2 (9.52) |
| APOE4 (%) | 465 (21.99) | 111 (19.65) | 75 (20.55) | 185 (27.29) | 94 (18.54) | 71 (17.88) | 23 (18.55) | 7 (12.28) | 25 (22.32) | 16 (15.38) | 23 (19.49) | 9 (17.65) | 0 (0.00) | 14 (37.84) | 0 (0.00) |
| TMT (SD) | 96.22 (55.91) | 89.33 (62.94) | 91.55 (53.65) | 102.19 (43.03) | 99.26 (63.01) | 97.28 (52.80) | 83.27 (45.83) | 97.47 (55.93) | 112.89 (49.09) | 97.09 (58.41) | 92.09 (50.06) | 88.12 (52.22) | 82.44 (30.67) | 115.78 (53.46) | 64.14 (19.55) |

N = Sample Size Female (%) = Sample size of female participants (percentage) colledge (%) = Sample size of participants with some college education (percentage) age (mean ± SD) = Mean and standard deviation of age in years AD (%) = Sample size of participants with Alzheimer's Disease (percentage) MCI (%) = Sample size of participants with Mild Cognitive Impairment (percentage) APOE ε2 (%) = Sample size of participants carrying APOE ε2 allele (percentage) APOE ε4 (%) = Sample size of participants carrying APOE ε4 allele (percentage) TMT = Mean and standard deviation of Trail Making Test Parts B in seconds ALL = All samples Boston = Samples from Boston Center Columbia = Samples from Columbia Center Denmark = Samples from Denmark Center Pittsburgh = Samples from Pittsburgh Center

### Whole genome sequencing (WGS)

The McDonnell Genome Institute (MGI) at Washington University in Saint Louis performed 150bp paired-end reads whole genome sequencing (WGS) using an Illumina sequencer, aligned reads to the Genome Reference Consortium Human Build 38 (GRCh38) with Burrows-Wheeler Aligner (BWA-MEM 0.7.15)(20), marked duplicates using Picard 2.4.1 (http://broadinstitute.github.io/picard/), recalibrated base quality score with GATK BaseRecalibrator 3.6(21), and converted to CRAM format with SAMtools 1.3.1 (http://www.htslib.org/doc/1.3.1/samtools.html). Variant calling was completed using the following four steps on the CRAM files: 1) The subject-level GVCF files were generated from CRAM files using GATK HaplotypeCaller(21); 2) All GVCF files were combined into one file with GATK CombineGVCF(21); 3) Genotypes were identified with GATK GenotypeGVCFs(21); and 4) The diallelic SNPs and INDELs were extracted to VCF files using GATK SelectVariants(21). Samples and variants that had low quality as defined by the following criteria: freemix contamination >3%, haploid coverage < 20×, and high Mendelian errors (the threshold of Mendelian errors differed by MAF(22)) reported by LOKI 2.4.5 (https://sites.stat.washington.edu/thompson/Genepi/Loki.shtml) and KING (Kinship-based Inference for GWAS; https://www.kingrelatedness.com/) version 2.3.1 were removed. Additional problematic variant sites determined by read depth < 20 or > 300, call rate < 90%, and excess heterozygosity (P_HWE_ < 1×10^-6^) were also removed from the analysis. A final comparison of the resultant WGS against previous SNP chip data was performed and led to the correction of a few sample swaps. Finally, we used Eigenstrat(23) on the cleaned WGS data to estimate principal components (PCs)(22).

### RNA sequencing (RNA-Seq)

MGI utilized the Qiagen QIAcube155 extraction robot to extract the total RNA from PAXgene^TM^ Blood RNA tubes with the Qiagen PreAnalytiX PAXgene Blood miRNA Kit (Qiagen, Valencia, CA) and conducted the whole genome paired-end RNA sequencing. The Division of Computational & Data Sciences at Washington University processed the RNA sequencing data using nf-core/rnaseq 3.14.0 (https://nf-co.re/rnaseq/3.14.0)(24). The major steps of processing involve alignment of the reads to GRCh38 with STAR (25), marking read duplicates with Picard MarkDuplicates (http://broadinstitute.github.io/picard/), and transcript assembly and quantification with StringTie(26). Poor quality genes, those with fewer than three counts per million in at least 98.5% of samples were dropped. Samples with more than 8% of reads mapped to intergenic regions were also excluded. The gene expression of cleaned data was transformed using the variance stabilizing transform (VST) function in DESeq(27), followed by estimation of principalcomponents using factoextra R package (https://cran.r-project.org/web/packages/factoextra/index.html), and covariate adjustment forcing age, age squared, sex, field center, percent of reads mapping to intergenic sequence, and the counts of red blood cells, white blood cells, platelets, monocytes, and neutrophils, and then stepwise regressing out the influences of RNA-seq batch and the top 10 PCs of gene expression. The adjusted gene expression residuals were used in our analyses.

### Lipidomics assay

The Biomedical Mass Spectrometry Lab at Washington University in Saint Louis carried out untargeted metabolomics workflow via liquid chromatography/mass spectrometry (LC/MS)(28, 29). Methyl tert-butyl ether/methanol was utilized to extract lipid metabolites from plasma samples in a solid-phase extraction (SPE) plate. Positive reverse-phase (RP) chromatography coupled to high-resolution mass spectrometry (HRMS) was conducted to obtain the m/z values of the extracted metabolites. Lipid iterative MS/MS data was annotated with the Agilent Lipid Annotator software (Agilent Technologies). The peak areas of the annotated peak list were estimated using Skyline (Version 20.1.0.155). Missing values were replaced by half of the minimum value. Using a pooled quality control sample as an internal standard, batch effects were corrected by a random forest based method(30). After QC, 188 lipids from 13 compound classes remained and were used for analysis. We further log2 transformed the peak area of each lipid to approximate normal distribution. The transformed lipid values were adjusted by age, sex, field centers, the top 10 PCs.

### Statistical Analyses

#### Phenotype

The time in seconds, needed to complete TMT-B (N=2119) was obtained as part of the LLFS Visit 2. To approximate a normal distribution and control for confounding, we performed a natural logarithm transformation on TMT-B, and adjusted by age, age squared, field centers, sex, education, and 10 genetic PCs using SAS 9.4.

#### Genome-Wide Linkage Analyses (GWLS) and statistical fine-mapping of the linkage peak

To estimate the contribution of the shared genetics within pedigrees to the TMT-B scores, we computed multipoint IBD matrices with each of the pedigree using LOKI 2.4.5 (https://sites.stat.washington.edu/thompson/Genepi/Loki.shtml)(22, 31), and conducted GWLS using Sequential Oligogenic Linkage Analysis Routines (SOLAR Eclipse version 8.3.1)(32). To obtain multipoint IBD estimates, within each of 0.5 cM intervals, we selected up to five tightly linked SNPs and constructed haplotype using ZAPLO(33). The haplotypes were then used to generate multipoint IBD matrices. A significant linkage peak has LOD score above 3 in a genetic region. Since not all families in LLFS contributed to the linkage signal, we identified the set of families with pedigree-specific LOD scores exceeding 0.1 as linked families, and the top linked families as those families whose ranked pedigree-specific LOD scores summed to ≥ 6. We performed association analyses of the WGS variants (diallelic SNPs and INDELs) under the significant linkage region (the region with 1 LOD score drop from the linkage peak) with the TMT-B score separately for samples from linked families or top linked families. To determine the contribution of each of the sequenced variants to the linkage peak, the sequenced variants that had P<1×10^-4^ were included as a covariate and the LOD score was re-estimated using SOLAR separately for samples from linked families or top linked families. The candidate variants under the linkage peak were nominated if they met all of the following criteria: 1) Showed a P-value < 1×10^-4^ in at least one of two sample sets (linked families or top linked families); 2) Demonstrated an association P-value < 0.05 in two sample sets (linked families, and top linked families); and 3) Reduced the LOD score by more than 0.5 in both linked families and top linked families. To assess the collective impact of the multiple variants (SNPs and INDELs) under the linkage peak, we conducted a stepwise regression using SAS 9.4 with a significance threshold for entering variable sets at P<0.1 and for staying variables at P<0.05 separately for linked families and top linked families. Subsequently, we re-estimated the LOD score using SOLAR accounting for the selected variants.

#### GWLS sensitivity analyses

Due to the strong correlation between TMT-B and TMT-A (reported correlation coefficient=0.62)(16), TMT-B and cognitive status, and TMT-B and APOE haplotypes(34), we assessed if the linkage peaks for TMT-B might be influenced by these traits. To determine the impact of cognitive status, and APOE haplotypes on the linkage peaks of TMT-B, we conducted GWLS conditioning on each of TMT-B, cognitive status (Alzheimer’s Disease status: 0 for no and 1 for yes; Mild Cognitive Impairment: 0 for no and 1 for yes), and APOE haplotypes (APOE _ε_2 dosage: 0 for no _ε_2 allele, 1 for one copy of _ε_2 allele, and 2 for two copies of _ε_2 allele; APOE _ε_4 dosage: 0 for no _ε_4 allele, 1 for one copy of _ε_4 allele, and 2 for two copies of _ε_4 allele) and assessed the LOD score alterations.

#### Expression quantitative trait loci (eQTLs) analysis

To map the identified variants to the gene expression, we tested the association of variant dosage with each of the adjusted RNA-Seq residuals using a linear mixed model implemented in “kinship” and “lmekin” R package.

#### Metabolite quantitative trait loci (mQTLs) analysis

To test whether any lipidomic molecules regulated by the identified variants, the association of variant dosage with each of the adjusted lipid residuals was performed with a linear mixed model implemented in “kinship” and “lmekin” R package.

#### Bioinformatics annotation

The rsID number, and location of the variants related to genes were annotated with the Ensemble Variant Effect Predictor (VEP) release 107 (GRCh38.p13 assembly for Homo_sapiens)(35) and OASIS (https://edn.som.umaryland.edu/OASIS/LLFS/).The allele frequency of the variants in LLFS was compared with the allele frequency for European from the Allele Frequency Aggregator (ALFA) dataset (release version: 20230706150541) which was used as population reference. Additional evidence of eQTLs or pQTLs from brain or blood might provide crucial biological insights for our identified variants. As such, we examined eQTL from MetaBrain(36), Genotype-Tissue Expression (GTEx) Analysis V8(37), blood eQTLGen(38), as well as CSF pQTL(39).

## Results

### Characteristics of participants

As depicted in Table 1, across the four field centers, the Denmark field center had the highest number of participants for TMT-B while the US field centers were lower. We consistently noted a slightly higher percentage of females (TMT-B: 53%) in all LLFS samples, as well as across four field centers (>51% for TMT-B). The average age of 2115 samples is 69.97 years old (ranging from 45 to 103 across field centers). The majority of participants (>80%) from all field centers had attained some college education Except for the Denmark field center, the Alzheimer’s Disease (AD) status of three US centers was determined by a dementia review committee including neurologists and neuropsychologists who reviewed comprehensive cognitive assessments, interviews, and ratings of everyday function via the Clinical Dementia Rating (CDR) Scale. Despite mean age being older than 65 years old, samples included in our analyses exhibited a relatively low prevalence of AD (Table 1: 3.55% vs 10% among non-Hispanic Whites)(40) and Mild Cognitive Impairment (MCI) (Table 1: 7.93% vs 14.9% in people age 65 and older)(41). Corresponding to the lower rates of AD and MCI, we observed a relatively higher percentage of carriers (∼17% vs ∼11.1% in White by Belloy et al)(42) for the protective APOE ε2, and relatively lower percentage of carriers (∼22% vs ∼40.6% in White by Belloy et al)(42) for the risk associated APOE ε4 allele (Table 1). Additionally, we observed significantly higher values for the TMT-B in the Denmark field center (102.19 seconds) than the US field centers. The demographic information for linked family samples (N=397) and top linked family samples (N=118) is presented in Table 1.

### GWLS detected a significant linkage peak at chromosome 15

Our GWLS, estimated using SOLAR, revealed significant linkage at 15q25 (LOD=5.2084). After adjusting the TMT-B by TMT-A, the LOD score remained above 2.5 for 15q25 (LOD=2.77; Figure S1). This linkage peak was not influenced by APOE _ε_2 status, APOE _ε_4 status, nor cognitive status (Figure S2, S3, and S4). Not all LLFS families contributed to the identified linkage peak. To maximize our power for fine-mapping, we identified 76 linked families with pedigree-specific LOD scores exceeding 0.1, and 11 top linked families where the sum of the ranked pedigree-specific LOD scores exceeded 6. As depicted in Figure 1, utilizing the selected families, the LOD score exceeds 16 for 76 linked families, and exceeds 6 for top linked families. Using 1 LOD score drop support interval, we were able to narrow down the linkage peak on chromosome to an ∼8 Mbp region (80.8 Mbp - 88.2 Mbp) which encodes 41 proteins.

**Figure 1.**
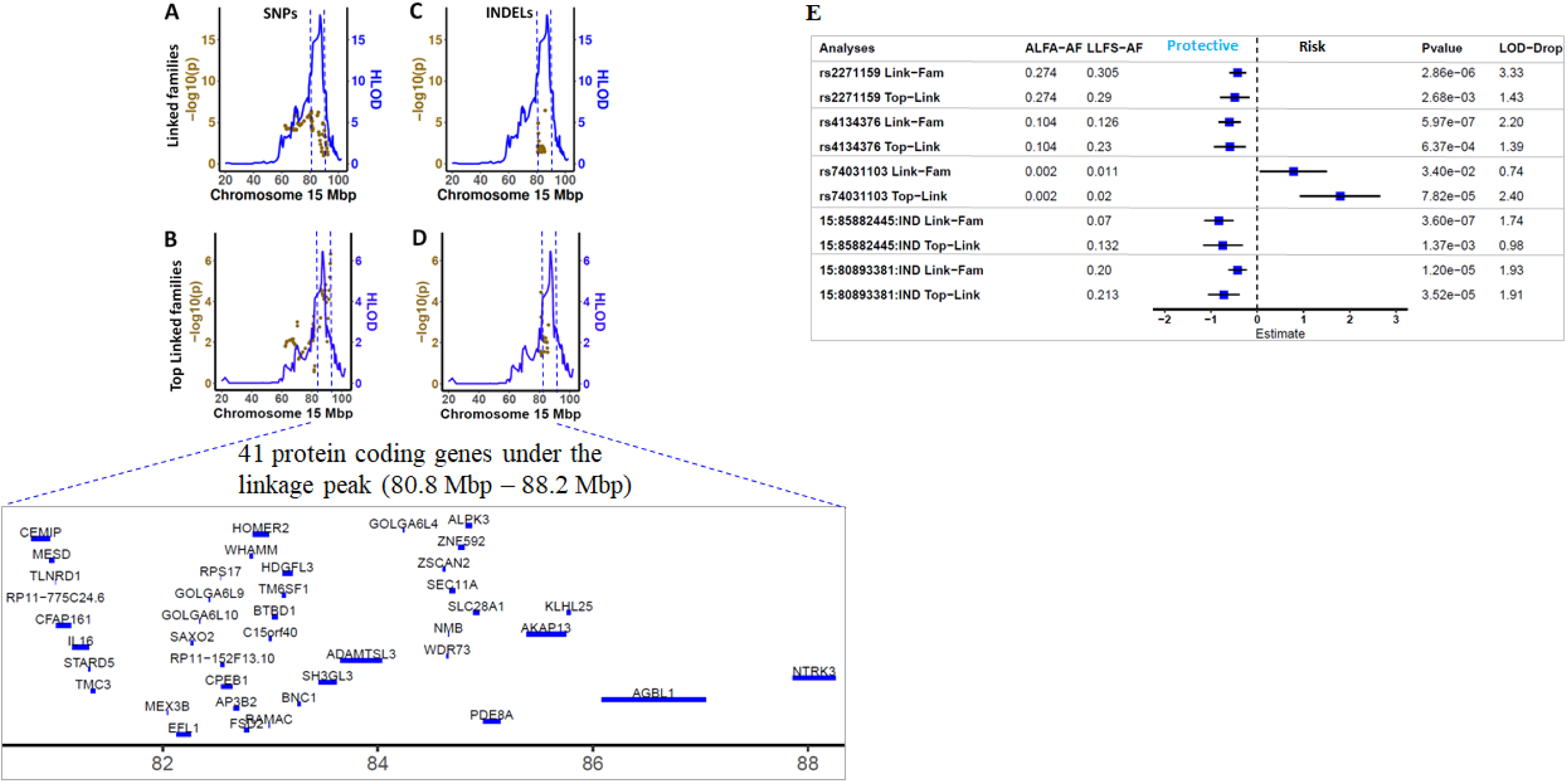
Linkage and association analyses for TMT-B at chromosome 15. A) Linkage and association analyses of SNPs at chromosome 15 using linked families. The x-axis is the physical distance in Mbp at chromosome 15. The y-axis (left) denotes the negative of log10 of two-sided P value, and the y-axis (right) indicates the B) Linkage and association analyses of SNPs at chromosome 15 using top linked families. The x-axis is the physical distance in Mbp at chromosome 15. The y-axis (left) denotes the negative of log10 of two-sided P value, and the y-axis (right) indicates the HLOD score of the linkage analyses. C) Linkage and association analyses of INDELs at chromosome 15 using linked families. The x-axis is the physical distance in Mbp at chromosome 15. The y-axis (left) denotes the negative of log10 of two-sided P value, and the y-axis (right) indicates the D) Linkage and association analyses of INDELs at chromosome 15 using top linked families. The x-axis is the physical distance in Mbp at chromosome 15. The y-axis (left) denotes the negative of log10 of two-sided P value, and the y-axis (right) indicates the HLOD score of the linkage analyses. E) Forest plots of five genetic variants contributing to the chromosome 15 peak in linked families and top linked families.

### Five genetic variants contributed to the linkage peak at chromosome 15

Under this peak, we identified five variants (three diallelic SNPs and two INDELs; Table 2 and Figure 1E) influencing the linkage peak by: 1) having P<1×10^-4^ in at least one of two sample sets (linked families or top linked families); 2) showing association P < 0.05 in all two sample sets (linked families, and top linked families); and 3) lowering the LOD score > 0.5 in both linked families and top linked families. Four of these five variants are protective (*CEMIP*-rs2271159, rs4134376, *KLHL25*-15:85882445:IND, and *CEMIP*-15:80893381:IND), showing a shorter completion time of TMT-B for carriers.

**Table 2.** Associations of diallelic SNPs under the linkage peak for TMT-B.

| rsID | CHR | BP | SYMBOL | Consequence | alfa_af | EA | NonEA | Linked Families |  |  |  |  |  | Top Linked Families |  |  |  |  |  |
| --- | --- | --- | --- | --- | --- | --- | --- | --- | --- | --- | --- | --- | --- | --- | --- | --- | --- | --- | --- |
|  |  |  |  |  |  |  |  | af | n | beta | se | pval | lod_drop | af | n | beta | se | pval | lod_drop |
| rs2271159 | 15 | 80887585 | CEMIP | intron | 0.2743 | T | G | 0.305 | 393 | -0.42 | 0.09 | 2.86E-06 | 3.33 | 0.29 | 115 | -0.48 | 0.16 | 2.68E-03 | 1.43 |
| rs4134376 | 15 | 85885026 | - | intergenic | 0.104 | G | A | 0.126 | 393 | -0.6 | 0.12 | 5.97E-07 | 2.2 | 0.23 | 117 | -0.59 | 0.17 | 6.37E-04 | 1.39 |
| rs74031103 | 15 | 88208591 | NTRK3 | intron | 0.0021 | C | A | 0.011 | 392 | 0.78 | 0.37 | 3.40E-02 | 0.74 | 0.02 | 117 | 1.79 | 0.44 | 7.82E-05 | 2.4 |
|  | 15 | 85882445 | KLHL25 |  |  | C | CTT | 0.07 | 379 | -0.83 | 0.16 | 3.60E-07 | 1.74 | 0.132 | 110 | -0.74 | 0.22 | 1.37E-03 | 0.98 |
|  | 15 | 80893381 | CEMIP |  |  | A | AG | 0.2 | 378 | -0.43 | 0.1 | 1.20E-05 | 1.93 | 0.213 | 108 | -0.72 | 0.17 | 3.52E-05 | 1.91 |

rsID = Reference SNP cluster ID CHR = Chromosome number BP = Physical distance in base pairs SYMBOL = gene name Consequence = Variant type alfa\_af = Allele Frequency in ALFA EUR EA = Effect Allele NonEA = Noneffect Allele af = Allele Frequency n = sample size beta = effect size estimate se = standard error Pval = two-sided P value lod\_drop = LOD score drop (lod\_pre - lod\_post)

As depicted in Figure 1E and Table 2, the first protective common intronic variant of *CEMIP*, rs2271159 (MAF ∼29% in LLFS), displays a significant protective effect in linked families (β=-0.42,P=2.86×10^-6^; Table 2) and top linked families (β=-0.48,P=2.69×10^-3^; Table 2), and explains 18% of the linkage peak in linked families, and 22% of the linkage peak in top linked families. To gain insights of function of this variant, we queried the genes regulated by this variant in MetaBrain eQTL (brain tissue), GTEx eQTL (brain tissue), eQTLGen (blood), LLFS eQTL (blood), as well as CSF pQTL. This variant is a strong cis-eQTL of *CEMIP* in the cerebellum (MetaBrain eQTL: β=0.597,P=4.49×10^-19^; Figure 2A), and a weak cis-eQTL of *CEMIP* in the Hippocampus (MetaBrain eQTL: β=0.303,P=1.25×10^-2^; Figure 2A). We did not observe the influences of rs2271159 on *CEMIP* in other brain regions, nor in blood. Interestingly, rs2271159 is weak trans-pQTLs of several proteins (TJP1, EMC4, GOLM2, SORD, COPS2, HERC1, and UBE2Q2; Figure 2B) in CSF. All these proteins were encoded by region outside of the linkage peak on Chromosome 15. Querying Open Targets identified the association of TJP1 with cortical surface area(43), EMC4 with cognitive ability (44), GOLM2 with neuritic plaque (45), SORD with cortical thickness (46), COPS2 with cortical thickness (46), HERC1 with Alzheimer’s disease (47), and UBE2Q2 with cortical surface area (43). This indicates that these proteins might be the molecules mediating the effects of rs2271159 on TMT-B. In addition, SM(d42:3) [Sphingomyelin] was also modulated by rs2271159 (Figure 2B). The involvement of Sphingomyelin metabolism in Alzheimer’s disease(48) supports SM(d42:3) might be a candidate that links rs2271159 to TMT-B.

**Figure 2.**
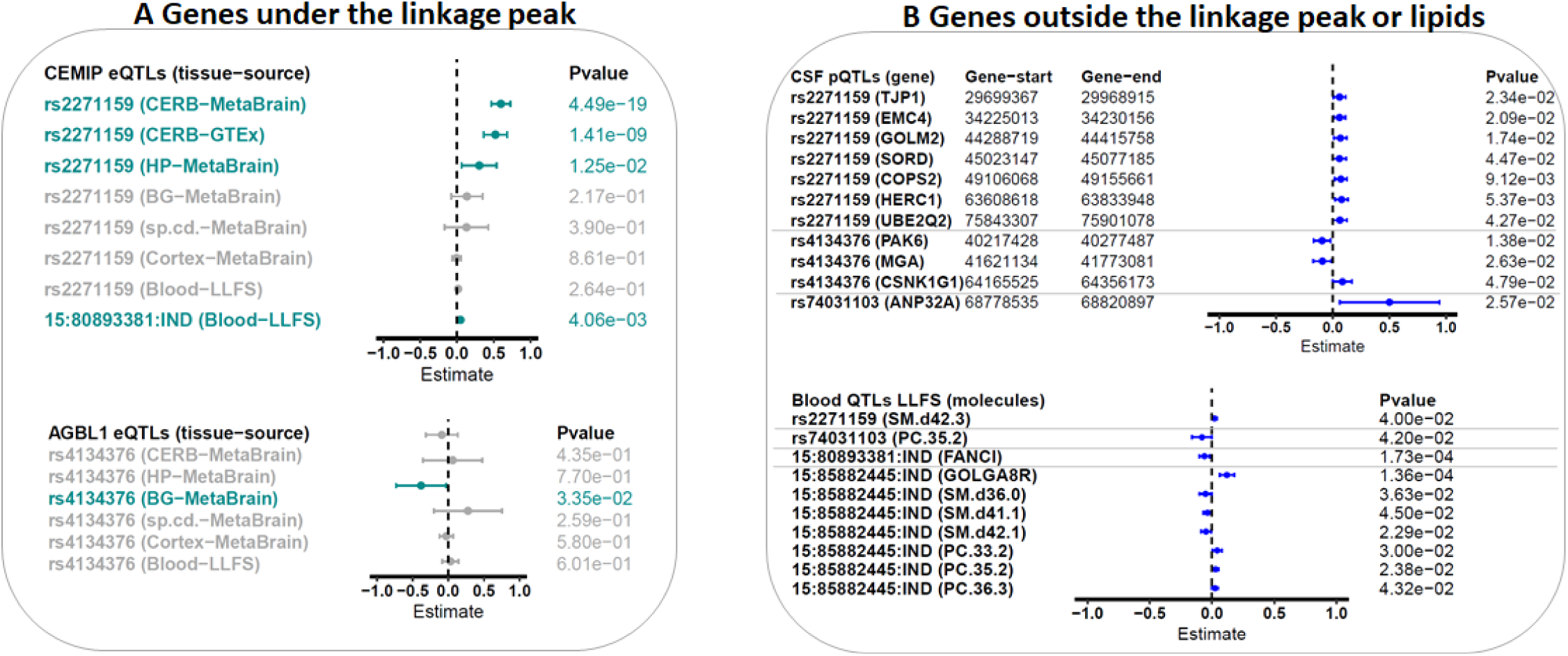
Forest plots of variants’ effects on genes under the linkage peak (A) or genes outside the linkage peak or on lipids (B) A) Forest plots of variants’ effects on CEMIP and AGBL1 under the linkage peak in blood as well as various brain regions. B) Forest plots of variants’ effects on genes residing outside the linkage peak in CSF and blood, as well as on lipids.

The second protective common variant, rs4134376 (MAF 12-23% in LLFS), showed minor allele enrichment in LLFS compared to the ALFA European database (MAF=10%), and explained 12% of the linkage in linked families, and 20% of the linkage in top linked families. This variant is associated with a protective effect on TMT-B (decreasing the time to complete the task) which was consistent across both sets of families analyzed (linked families, β=-0.6,P=5.97×10^-7^; and top linked families, β=-0.59,P=6.40×10^-4^; Table 2). A weak association between rs4134376 and *AGBL1* was noted specifically in basal ganglia (MetaBrain eQTL: P=0.03; Figure 2A)(36), as well as having weak associations with several CSF proteins (CSF pQTL: PAK6, MGA, and CSNK1G1)(39). *AGBL1* is a learning performance related gene(49), PAK6 is related to cognitive performance (50), MGA is associated with cortical thickness (46), and CSNK1G1 influences CSF t-tau . These results support biologic plausibility of these genes’ involvement with the TMT-B task (51).

Furthermore, we identified two common INDELs (15:80893381 within *CEMIP* and 15:85882445 downstream of *KLHL25*) which were protective for TMT-B across both sample sets (P<0.05) and contributed to ∼11% of the linkage peak in linked families and >19% of the linkage peak in the top linked families. The INDEL 15:80893381 is a cis-eQTL of *CEMIP* (MetaBrain eQTL: P=4.06×10^-3^; Figure 2A) and a trans-eQTL of *FANCI* in blood using LLFS samples (P=1.73×10^-4^; Figure 2B). The INDEL 15:85882445 is an eQTL for *GOLGA8R* in blood using LLFS samples (P=1.36×10^-4^; Figure 2B). The association of *FANCI* with white matter microstructure (52), and *GOLGA8R* locus with cortical surface area(43, 46) supports that these genes might link the INDELs to cognitive function. In addition, multiple metabolites (SM(d36:0) [Sphingomyelin], SM(d41:1)[Sphingomyelin], SM(d42:1)[Sphingomyelin], PC 33:2 [Phosphatidylcholine], PC 35:2 [Phosphatidylcholine], and PC 36:3 [Phosphatidylcholine]) were nominally associated with 15:85882445 (P<0.05).

The final variant was a rare intronic variant of *NTRK3*, rs74031103, with significantly enriched minor allele frequency in LLFS (MAF=1%-2%) compared to the ALFA European database (MAF=0.2%). This SNP is significantly associated with longer completion time on the TMT-B (ie harmful) (β=1.79,P=7.82×10^-5^; Table 2 and Figure 1E), and explained 37% of linkage in the top linked families. The association was consistent in the linked families (β=0.78,P=3.40×10^-2^; Table 2). Two molecules (CSF ANP32A and blood PC 36:3 [Phosphatidylcholine]) were regulated by rs74031103 (P<0.05; Figure 2B).

### Influence of the linkage peak jointly by diallelic SNPs and INDELS

We identified a total of five variants (three SNPs and two INDELs) under the chromosome 15 linkage peak for TMT-B. Except for the moderate Linkage Disequilibrium (LD) observed between rs2271159 and 15:80893381 (r^2^=0.56), and between rs4134376 and 15:85882445 (r^2^=0.53), LD between the remaining variant pairs is negligible (r^2^ 0). To assess the impact of the variants collectively, we selected the significant independent variants using a stepwise regression with entering threshold at P<0.1 and staying threshold at P<0.05. We performed the selection procedure separately for each of two sets of samples, and selected three variants for linked families, as well as for top linked families. The SNP rs4134376 was selected for both sample sets. The remaining two variants selected were different for linked families (rs2271159 and 15:85882445) and top linked families (rs74031103 and 15:80893381), indicating that strength of the variants differed by family sets. The selected variants dropped the LOD score from 16.08 to 11.27, and explained ∼30% of the linkage peak for linked families (Figure 3). Notably, the linkage peak for top linked families disappeared (LOD score dropped from 6.02 to 1.46) after adjusting the three selected variants.

**Figure 3.**
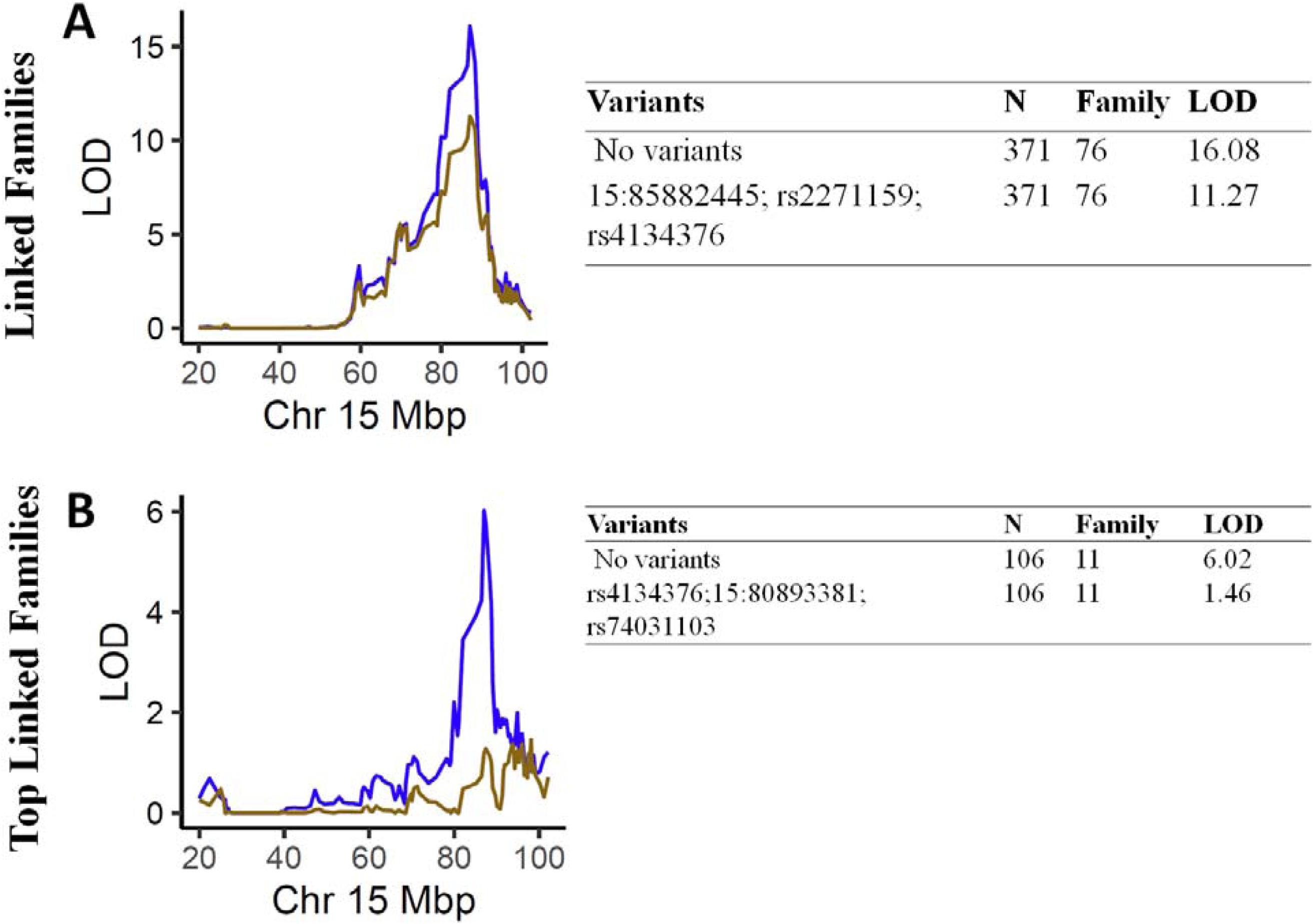
Linkage analyses for TMT-B before and after adjusting for SNPs and INDELs. A) Plots of linkage results for TMT-B using linked family for chromosome 15. The x-axis denotes the physical distance in megabase pair (Mbp) at chromosome 15. The y-axis indicates the LOD Score of the linkage analyses. The blue (chr 15) curve depicts the linkage results before adjusting for SNPs and INDELs. The brown curve denotes the linkage results after adjusting for SNPs and INDELs. B) Plots of linkage results for TMT-B using top linked family for chromosome 15. The x-axis denotes the physical distance in megabase pair (Mbp) at chromosome 15. The y-axis indicates the LOD Score of the linkage analyses. The blue (chr 15) curve depicts the linkage results before adjusting for SNPs and INDELs. The brown curve denotes the linkage results after adjusting for SNPs and INDELs.

## Discussion

We conducted GWLS on TMT-B performance using a cohort of over 2000 participants from long-lived families. We document significant linkage at 15q25. Our fine mapping under 15q25 detected five variants (three SNPs: *NTRK3*-rs74031103, *CEMIP*-rs2271159, and rs4134376; two INDELs: *KLHL25*-15:85882445, *CEMIP*-15:80893381) for TMT-B. A rare intronic risk SNP rs74031103 in *NTRK3* was noted for TMT-B. More importantly, four of the variants (*CEMIP*-rs2271159, rs4134376, *KLHL25*-15:85882445, *CEMIP*-15:80893381) had protective effects on TMT-B with their minor alleles. Notably, one SNP (rs2271159) and one INDEL (15:80893381) reside in *CEMIP*. Although the association of rs2271159 with TMT-B did not reach nominal significance using all LLFS families, the consistent negative effect of its T allele was observed in all LLFS families, linked families, and top-linked families. Additionally, elevated levels of *CEMIP* in the cerebellum and hippocampus were significantly associated with its T alleles, along with its known involvement in maintaining the central nervous system (CNS).

*CEMIP* is known to contribute to the maintenance and homeostasis of the CNS by facilitating the degradation of hyaluronic acid (HA), a primary component of the brain’s extracellular matrix(53). *CEMIP* is the only hyaluronidase expressed in brain that depolymerize hyaluronic acid into small-and intermediate-sized fragments. The involvement of hyaluronic acid in regulation of synapse function and decreased mnemonic ability in *CEMIP* knock-out mice (54) support that the protective effect of rs2271159 was mediated through enhanced *CEMIP* expression in cerebellum and hippocampus. Additionally, several molecules including seven CSF proteins (TJP1, EMC4, GOLM2, SORD, COPS2, HERC1, and UBE2Q2) and one lipid (SM(d42:3) [Sphingomyelin]) were also influenced by this SNP. Reported associations with cortical structure (43), cognitive ability (44), neuritic plaques (45), and Alzheimer’s disease (47) (48) for these molecules indicates that they might be also involved in modulating cognitive function as assessed by TMT-B.

The second protective common variant, rs4134376, explained 12% - 20% of the linkage peak. This SNP is a basal ganglia specific eQTL of *AGBL1* where a lower level of AGBL1 in basal ganglia was associated with its G allele. AGBL1 encodes a glutamate decarboxylase that mediates deglutamylation of tubulin.

Combining the facts that glutamylation stabilizes microtubules, higher AGBL1 expression may be detrimental to brain function. Therefore, the protective effect of rs4134376 is likely through lowering AGBL1 expression in basal ganglia. In addition, the three proteins (PAK6, MGA, and CSNK1G1) were also influenced by rs4134376 and may also be plausible candidate molecules. *AGBL1* is a learning performance related gene(49), PAK6 is related to cognitive performance(50), MGA is associated with cortical thickness(46), and CSNK1G1 influences CSF t-tau. Thus there is mounting evidence to support biologic plausibility of these genes’ involvement with the TMT-B task(51).

*CEMIP* and *FANCI* are genes that mediated the effect of INDEL 15:80893381, and *GOLGA8R* is the gene influenced by INDEL 15:85882445. The association of *FANCI* with white matter microstructure (52), and *GOLGA8R* locus with cortical surface area(43, 46) supports that these genes might link the INDELs to TMT-B. In addition, multiple metabolites (SM(d36:0) [Sphingomyelin], SM(d41:1)[Sphingomyelin], SM(d42:1)[Sphingomyelin], PC 33:2 [Phosphatidylcholine], PC 35:2 [Phosphatidylcholine], and PC 36:3 [Phosphatidylcholine]) were associated with 15:85882445 (P<0.05).

*NTRK3* and ANP32A are the potential functional molecules regulated by the rare intronic risk SNP rs74031103 in *NTRK3. NTRK3* encodes a member of the neurotrophic tyrosine receptor kinase (NTRK) family and is involved in the development of the neurons. Variants in ANP32A locus were associated with cortical surface area(43).

Finally, three variants (rs2271159, rs4134376, and 15:80893381) selected by a stepwise regression completely explained the linkage peak in top linked families. This is highly significant evidence that these three variants are either the variants responsible for the peak (i.e. explaining the better TMT-B time) or they are in high linkage disequilibrium with the acting variants. Whichever case it is, it signifies that the region identified on Chromosome 15 is highly correlated to TMT-B and thereby the various cognitive.

In this study, we integrated multi-omics data (WGS, transcriptomic data, and lipidomic data) and utilized various eQTL resources to gain a deeper understanding of the identified variants. To reduce the false positive rate, we conducted our linkage analyses using two sets of samples (linked families, and top linked families), thereby demonstrating reproducibility. More crucially, LLFS comprises exceptionally long-lived multi-generational families, offering a unique opportunity to discover novel genomic regions harboring variants for a trait of interest, in this case, TMT-B, within this context.

While our study presents novel findings and highlights its strengths, it is not immune to limitations. Firstly, our LLFS participants were selected from exceptionally long-lived families, thus our findings may not be readily generalizable to the broader population. Secondly, our analyses were conducted using samples of European descent, which may limit the generalizability of our results to other ancestral groups, such as African Americans and Asians. Thirdly, it’s important to note that our findings are based solely on statistical association analyses. Therefore, definitive conclusions regarding the causality of these variants and genes cannot be drawn without further experimental validations.

## Conclusion

Utilizing LLFS and multi-omic data, our GWLS identified novel genetic variants, mapped the functional molecules, and gained deeper insights into regulation of TMT-B performance.

## Supporting information

TrailsB_supplemental

## Ethics declarations

### Authors’ relationships and activities

The authors declare that there are no relationships or activities that might bias, or be perceived to bias their work.

### Contribution statement

LW, KT, MW analyzed the data. SLA and SC involved in data collection. VAM, EWD, JAA, SJL, and AS assisted in analyses and QC. MP and MW contributed to the conception and design of this manuscript. All authors approved the final version of the manuscript.

