## Supplementary material for "Identification of novel protective loci for executive function using the trail making test part B in the Long Life Family Study": TrailsB_supplemental

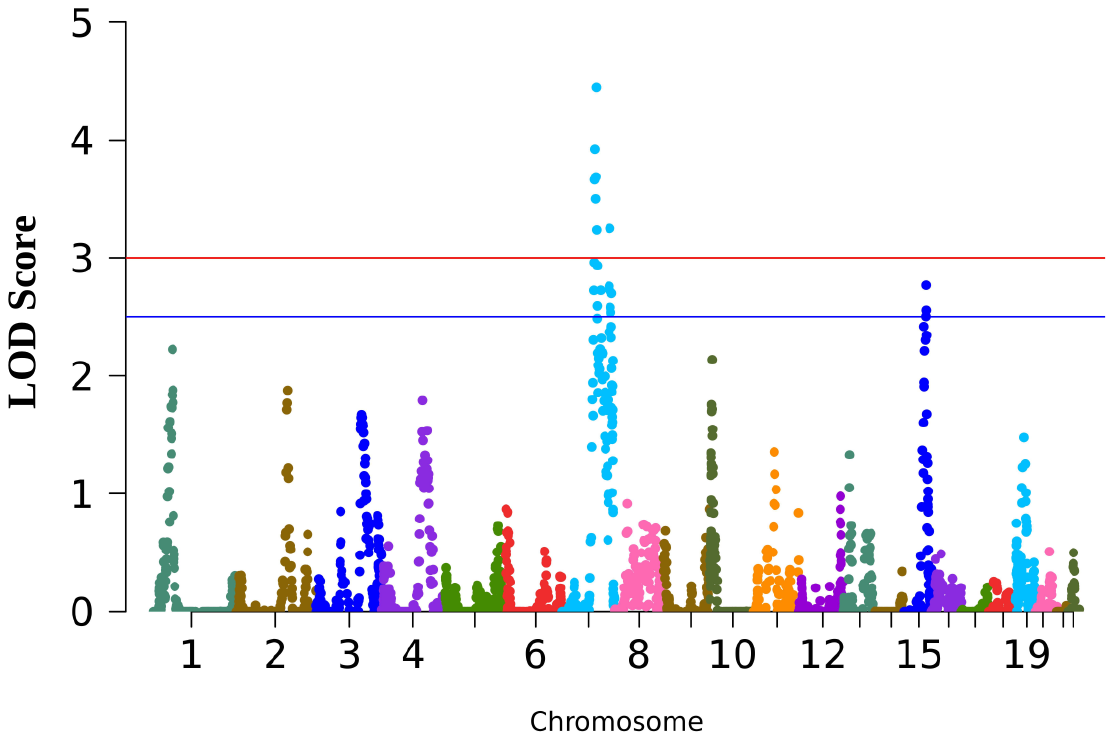

**Figure S1. GWLS of TMT-B adjusted by TMT-A**

Plots of linkage analyses across 22 chromosomes. The x-axis denotes the physical distance in base pairs by 22 chromosomes. The y-axis indicates the LOD Score of the linkage analyses. LOD score is 4.4451 at chromosome 7, 1.3552 at chromosome 11, and 2.7687 at chromosome 15.

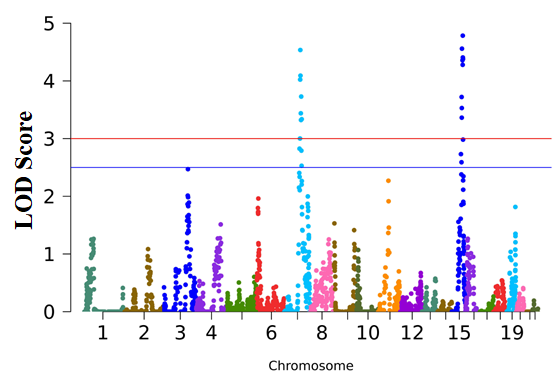

**Figure S2. GWLS of TMT-B adjusted by cognitive status**

Plots of linkage analyses across 22 chromosomes for TMT-B adjusted by dosage of cognitive status (Alzheimer’s Disease coded 1 for yes and 0 for no, and mild cognitive impairment coded 1 for yes and 0 for no). The x-axis denotes the physical distance in base pairs by 22 chromosomes. The y-axis indicates the LOD Score of the linkage analyses. LOD score is 4.5353 at chromosome 7, 2.2695 at chromosome 11, and 4.7841 at chromosome 15.

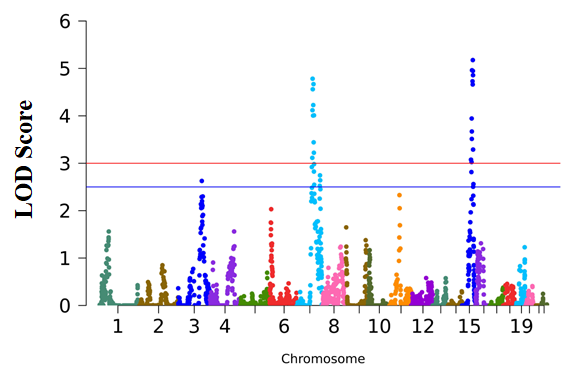

**Figure S3. GWLS of TMT-B adjusted by APOE**

Plots of linkage analyses across 22 chromosomes for TMT-B adjusted by dosage of APOE Ɛ2 (Ɛ2/Ɛ2 coded as 2, one copy of Ɛ2 coded as 1, and zero copy of Ɛ2 coded as 0) and dosage of APOE Ɛ4 (Ɛ4/Ɛ4 coded as 2, one copy of Ɛ4 coded as 1, and zero copy of Ɛ4 coded as 0). The x-axis denotes the physical distance in base pairs by 22 chromosomes. The y-axis indicates the LOD Score of the linkage analyses. LOD score is 4.7832 at chromosome 7, 2.3280 at chromosome 11, and 5.1762 at chromosome 15.

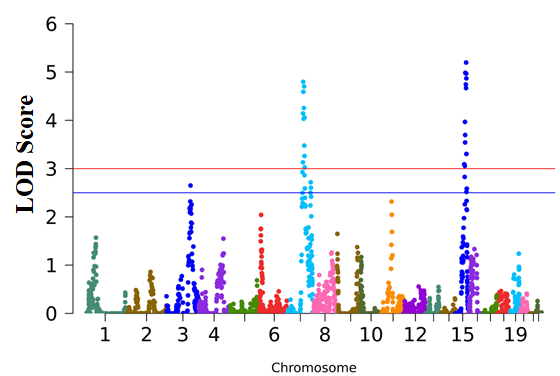

**Figure S4. GWLS of TMT-B adjusted by APOE**

Plots of linkage analyses across 22 chromosomes for TMT-B adjusted by dosage of APOE Ɛ2 (Ɛ2/Ɛ2 coded as 2, one copy of Ɛ2 coded as 1, and zero copy of Ɛ2 coded as 0). The x-axis denotes the physical distance in base pairs by 22 chromosomes. The y-axis indicates the LOD Score of the linkage analyses. LOD score is 4.7984 at chromosome 7, 2.3164 at chromosome 11, and 5.1971 at chromosome 15.

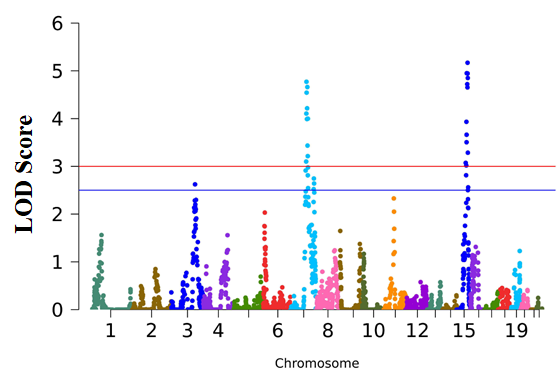

**Figure S5. GWLS of TMT-B adjusted by APOE**

Plots of linkage analyses across 22 chromosomes for TMT-B adjusted by dosage of APOE Ɛ4 (Ɛ4/Ɛ4 coded as 2, one copy of Ɛ4 coded as 1, and zero copy of Ɛ4 coded as 0). The x-axis denotes the physical distance in base pairs by 22 chromosomes. The y-axis indicates the LOD Score of the linkage analyses. LOD score is 4.7712 at chromosome 7, 2.3280 at chromosome 11, and 5.1697 at chromosome 15.
